# Modelling the effects of bacterial cell state and spatial location on tuberculosis treatment: Insights from a hybrid multiscale cellular automaton model

**DOI:** 10.1101/059113

**Authors:** Ruth Bowness, Mark A. J. Chaplain, Gibin G. Powathil, Stephen H. Gillespie

## Abstract

If improvements are to be made in tuberculosis (TB) treatment, an increased understanding of disease in the lung is needed. Studies have shown that bacteria in a less metabolically active state, associated with the presence of lipid bodies, are less susceptible to antibiotics, and recent results have highlighted the disparity in concentration of different compounds into lesions. Treatment success therefore depends critically on the responses of the individual bacteria that constitute the infection.

We propose a hybrid, individual-based approach that analyses spatio-temporal dynamics at the cellular level, linking the behaviour of individual bacteria and host cells with the macroscopic behaviour of the microenvironment. The individual elements (bacteria, macrophages and T cells) are modelled using cellular automaton (CA) rules, and the evolution of oxygen, drugs and chemokine dynamics are incorporated in order to study the effects of the microenvironment in the pathological lesion. We allow bacteria to switch states depending on oxygen concentration, which affects how they respond to treatment. This is the first multiscale model of its type to consider both oxygen-driven phenotypic switching of the *Mycobacterium tuberculosis* and antibiotic treatment. Using this model, we investigate the role of bacterial cell state and of initial bacterial location on treatment outcome. We demonstrate that when bacteria are located further away from blood vessels, less favourable outcomes are more likely, i.e. longer time before infection is contained/cleared, treatment failure or later relapse. We also show that in cases where bacteria remain at the end of simulations, the organisms tend to be slower-growing and are often located within granulomas, surrounded by caseous material.

## 1. Introduction

Although tuberculosis (TB) has long been both preventable and curable, a person dies from tuberculosis approximately every eighteen seconds (WHO Global Health Report 2011). Current treatment requires at least six months of multiple antibiotics to ensure complete cure and more effective drugs are urgently needed to shorten treatment. Recent clinical trials have not resulted in a shortening of therapy and there is a need to understand why these trials were unsuccessful and which new regimen should be chosen for testing in the costly long-term pivotal trial stage (Gillespie et al., 2014).

The current drug development pathway in tuberculosis is imperfect as standard preclinical methods may not capture the correct pharmacodynamics of the antibiotics. Using *in vitro* methods, it is difficult to accurately reproduce the natural physiological environment of *Mycobacterium tuberculosis* (*M. tuberculosis*) and the reliability of *in vivo* models may be limited in their ability to mimic human pathophysiology.

When *M. tuberculosis* bacteria enter the lungs, a complex immune response ensues. The outcome of infection is dependent on how effective the host’s immune system is and on the pathogenicity of the bacteria. The majority of patients will be able to control infection and contain it within a granuloma, which is a combination of immune cells that surround the bacteria. The centre of the granuloma may exhibit caseous necrosis and have a cheese-like appearance (Canetti et al., 1955). Most commonly, granulomas will undergo fibrosis or calcification and the infection is contained and becomes latent (Canetti et al., 1955; Grosset, 1980). In these cases, however, the individuals are still at risk of future relapse. If the granulomas do not contain the disease and infection continues, the bacteria can grow extracellularly.

Although a minimum of six months of therapy is recommended, it has long been recognised that many patients require less and are culture negative in two months or less (Fox, 1981). Shortening treatment to four months or less results in unacceptably high relapse rates (Gillespie et al., 2014) and studies (Study, 1981; Singapore, 1981) described in (Fox et al., 1999). It has recently been shown that, for some patients who become culture negative in only a week, there is still a non-zero risk of relapse (Phillips et al., 2016).

Figure 1 describes the estimates of tuberculosis containment versus progression to active disease in the general population. In patients with established disease, the outcome is perhaps determined by the ability of antibiotics to penetrate to the granuloma (Prideaux et al., 2015). In order to improve tuberculosis treatment, it is therefore vital to ensure sufficient concentrations of antibiotic reach the sites of infection (Via et al., 2015).

**Figure 1:**
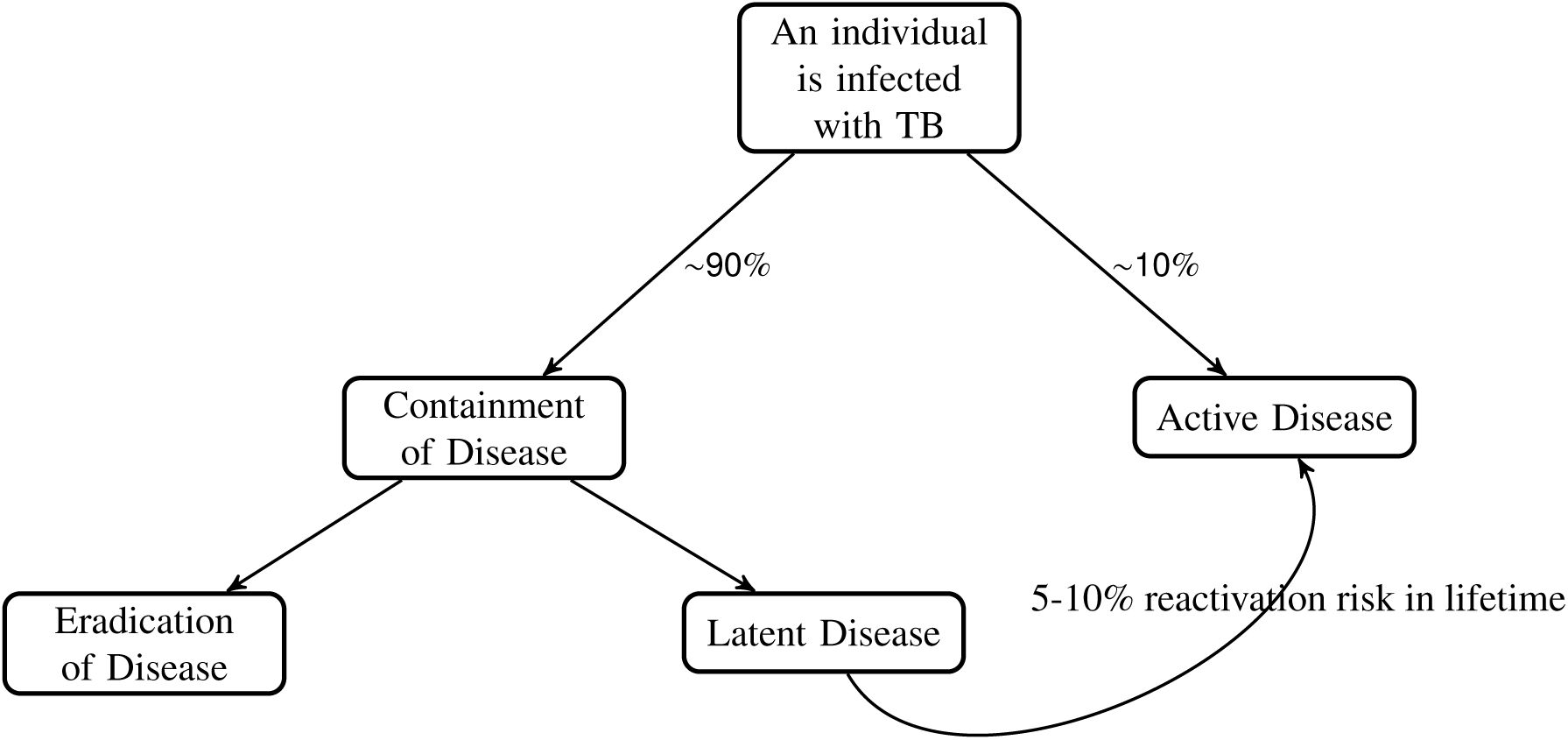
Schematic of population percentages of outcomes of TB disease. Data taken from (Ahmad, 2010) and references therein.

It is increasingly recognised that *M. tuberculosis* is able to enter into a state in which it is metabolically less active. Dormant bacteria have reduced susceptibility to antibiotics of which cell wall inhibitors are most affected but the action of the RNA polymerase inhibitor rifampicin and fluroquinolones acting on DNA gyrase is also reduced (Wayne and Hayes, 1996). A significant number of metabolic systems are down regulated in response to dormancy inducing stresses (Keren et al., 2004, 2011). This slower-growing state, associated with the presence of lipid bodies in the mycobacteria can increase resistance by 15 fold (Hammond et al., 2015). It has also recently been shown that around 60% of bacteria in the lung are lipid rich (Baron et al., 2017). The reduced susceptibility of some bacteria to antibiotics means that it is very important to study and analyse the heterogeneity of the bacteria so more effective treatment protocols can be developed. The spatial location of the bacteria is also vitally important as the ability of antibiotics to penetrate different sites of infection effectively is variable (Prideaux et al., 2015).

Multiple routes to dormancy have been reported and reviewed in detail (Lipworth et al., 2016). Oxygen concentration was one of the first mechanisms demonstrated in an *in vitro* model to result in dormancy, and *in vitro* models have been developed to explore the antibiotic susceptibility and metabolism of organisms in this slower-growing state (Wayne and Sramek, 1994). It has been hypothesised by multiple authors that lesions containing slower-growing bacteria are responsible for relapse (Grosset, 1980; Prideaux et al., 2015).

Cellular automaton modelling (and individual-based modelling in general) has been used to model other diseases, most notably tumour development and progression in cancer (Alarcón et al., 2003; Gerlee and Anderson, 2007; Zhang et al., 2009; Dormann and Deutsch, 2002; Swat et al., 2012; Powathil et al., 2012). Tuberculosis granulomas have been simulated previously through an agent-based model called ‘GranSim’ (Segovia-Juarez et al., 2004; Marino et al., 2011; Cilfone et al., 2013; Pienaar et al., 2015), which aims to reconstruct the immunological processes involved in the development of a granuloma.

In (Pienaar et al., 2016) the authors map metabolite and genescale perturbations, finding that slowly replicating phenotypes of *M. tuberculosis* preserve the bacterial population *in vivo* by continuously adapting to dynamic granuloma microenvironments. (Sershen et al., 2016) also combines a physiological model of oxygen dynamics, an agent-based model of cellular immune response and a systems-based model of *M.tb* metabolic dynamics. Their study suggests that the dynamics of granuloma organisation mediates oxygen availability and illustrates the immunological contribution of this structural host response to infection outcome.

In this paper, we build on previous models such as (Segovia-Juarez et al., 2004; Marino et al., 2011; Cilfone et al., 2013; Pienaar et al., 2015), and report the development of a hybrid-cellular automaton model. Our model is the first multiscale model to consider both oxygen dynamics and antibiotic treatment effects within a tuberculosis lesion, in order to investigate the role of bacterial cell state heterogeneity and bacterial position within the tuberculosis lesion on the outcome of disease. This is a unique focus for this type of model.

## 2. The hybrid multiscale mathematical model

The model simulates the interaction between TB bacteria, T cells and macrophages. Immune responses to the bacterial infection can lead to an accumulation of dead TB bacteria and macrophages, creating caseum. Oxygen diffuses into the system from blood vessels: bacteria switch from a slow-growing (lipid rich) to a fast-growing (lipid poor) phenotype in an oxygen-rich environment (proximate to the blood vessels). Chemokine molecules are secreted by the macrophages, which direct the movement of the immune cells. We then investigate the effect that antibiotics have on the infection.

Our spatial domain is a two dimensional computational grid, where each grid point represents either a TB bacterium, a macrophage, a T cell, caseum, the cross-section of a blood vessel or the extracellular matrix which goes to make up the local microenvironment. The spatial size of this computational grid has been chosen so that each automaton element is approximately the same size as the largest element in the system: the macrophage. At present we allow each grid cell to be occupied by a maximum of one element.

Our model is made up of five main components: (1) discrete elements - the grid cell is occupied either by a TB bacterium, a macrophage, a T cell, caseum or is empty. If the grid cell is occupied, automaton rules control the outcome; (2) the local oxygen concentration, whose evolution is modelled by a partial differential equation; (3) chemokine concentrations, modelled by a partial differential equation; (4) antibiotic concentrations, modelled by a partial differential equation and (5) blood vessels from where the oxygen and antibiotics are supplied within the domain. A schematic overview of the model is given in Figure 2.

**Figure 2:**
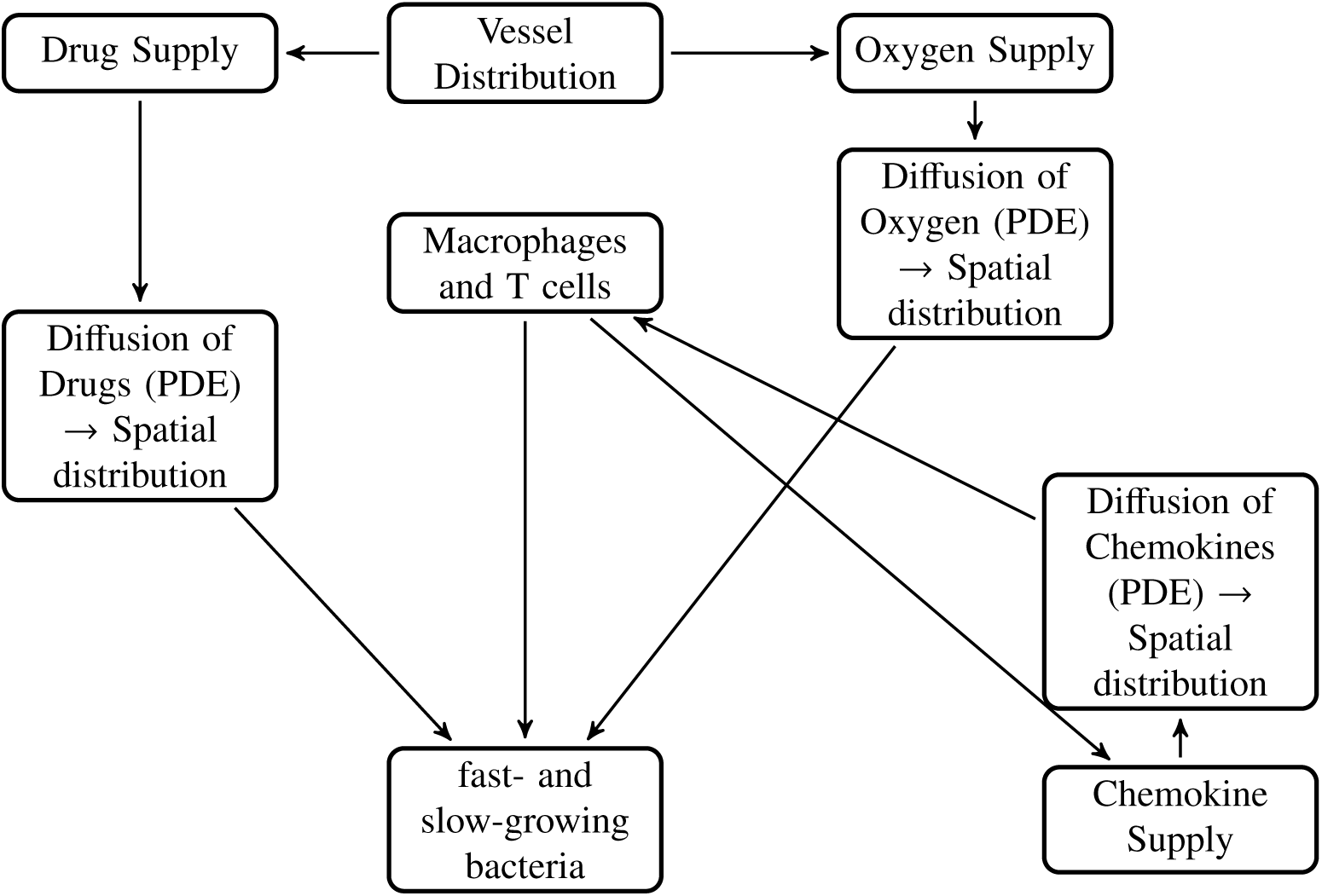
Schematic describing the basic processes in the model.

### 2.1 The blood vessel network

At the tissue scale, we consider oxygen and drug dynamics. We introduce a network of blood vessels in the model, which is then used as a source of oxygen and antibiotic within the model. Following (Powathil et al., 2012), we assume blood vessel cross sections are distributed throughout the two dimensional domain, with density *φ*_*d*_ = *N*_*v*_*/N*^2^, where *N*_*v*_ is the number of vessel cross sections (Figure 3). See Table 2 for values of *N* and *N*_*v*_. This is reasonable if we assume that the blood vessels are perpendicular to the cross section of interest and there are no branching points through the plane of interest (Patel et al., 2001; Daşu et al., 2003). We ignore any temporal dynamics or spatial changes of these vessels.

**Table 2:** Diffusion and decay parameters for antibiotics and Chemokine molecules

| Drug/Chemokine | Diffusion rate ($\text{cm}^2\text{s}^{-1}$) | Decay rate ($\text{hr}^{-1}$) |
| --- | --- | --- |
| Rifampicin (R) | $1.7 \times 10^{-6}$ | 0.17 |
| Isoniazid (H) | $1.5 \times 10^{-5}$ | 0.35 |
| Pyrazinamide (Z) | $1.6 \times 10^{-5}$ | 0.12 |
| Ethambutol (E) | $1.3 \times 10^{-5}$ | 0.2 |
| Chemokine | $10^{-6}$ | 0.347 |

**Figure 3:**
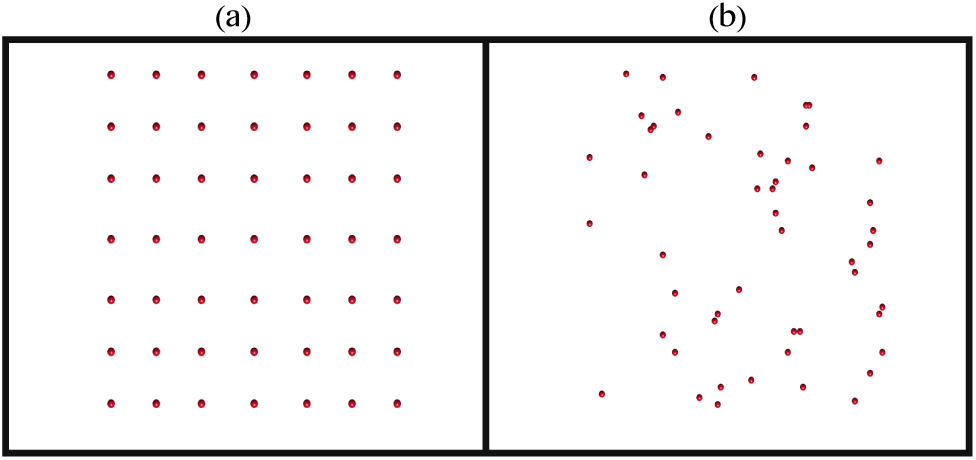
Plot illustrating (a) a fixed, uniform distribution of blood vessel cross sections throughout the spatial domain used in the cellular automaton simulations and (b) one outcome of a random distribution of the blood vessels.

### 2.2 Oxygen dynamics

The oxygen dynamics are modelled using a partial differential equation with the blood vessels as sources, forming a continuous distribution within the simulation domain. If *O*(**x**, *t*) denotes the oxygen concentration at position **x** at time *t*, then its rate of change can be expressed as

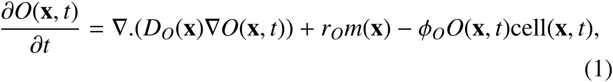

where *D*_*O*_(**x**) is the diffusion coefficient and *φ*_*O*_ is the rate of oxygen consumption by a bacterium or immune cell at position *x* at time *t*, with cell(**x**, *t*) = 1 if position *x* is occupied by a TB bacterium or immune cell at time *t* and zero otherwise. Oxygen consumption rates are different for each cell: 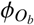 for the consumption by bacteria, 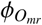 for resting macrophages, 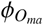 for active macrophages, 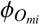 for infected macrophages, 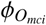 for chronically infected macrophages and 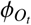 for T cells (see Table 2 for these values). *m*(**x**) denotes the vessel cross section at position **x**, with *m*(**x**) = 1 for the presence of blood vessel at position **x**, and zero otherwise; the term *r*_*O*_*m*(**x**) therefore describes the production of oxygen at rate *r*_*O*_. We assume that the oxygen is supplied through the blood vessel network, and then diffuses throughout the tissue within its diffusion limit. At this time, we assume a constant background oxygen level arising from airways and focus on oxygen diffusion from the vascular system. Since it has been observed that when a vessel is surrounded by caseous material, its perfusion and diffusion capabilities are impaired (Datta et al., 2015; Pienaar et al., 2016), we have incorporated this by considering a lower diffusion and supply rate in the granuloma structure as compared to the normal vessels, i.e.

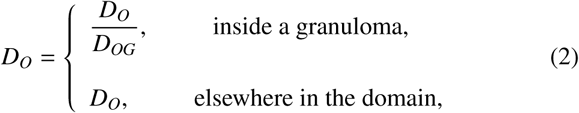

and

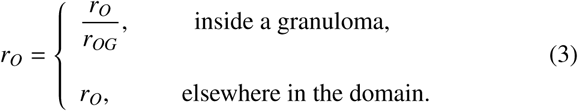

The formulation of the model is then completed by prescribing no-flux boundary conditions and an initial condition (Powathil et al., 2012). Figure 4 shows a representative profile of the spatial distribution of oxygen concentration after solving the Equation 1 with relevant parameters as discussed in Section 2.5.

**Figure 4:**
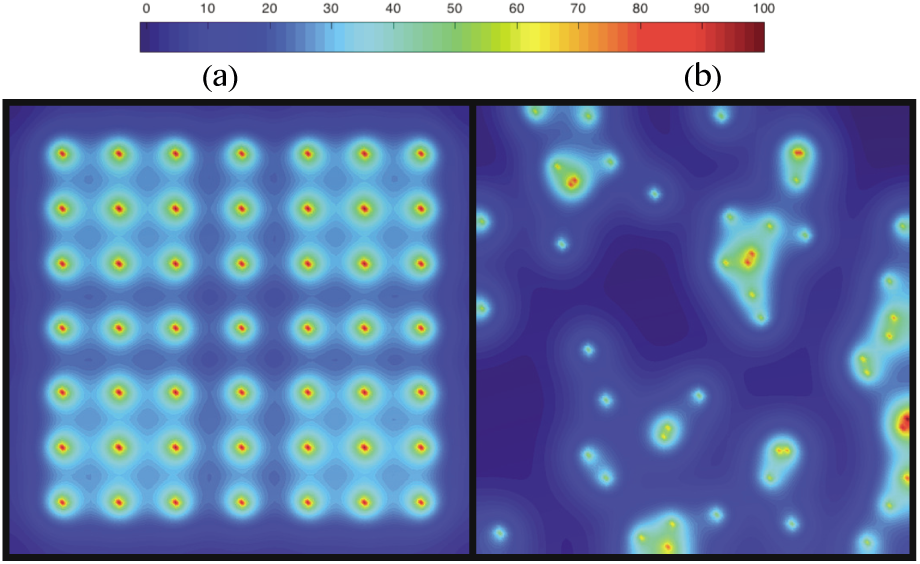
Plot showing the concentration profile of oxygen supplied from the blood vessel network for the (a) the fixed, uniform distribution of blood vessels shown in Figure 3 (a), and (b) the random distribution of blood vessels shown in Figure 3 (b). The red coloured spheres represent the blood vessel cross sections as shown in Figure 3 and the colour map shows the percentages of oxygen concentration.

### 2.3. Antibiotic treatments

In the present model we assume a maximum drug effect, allowing us to concentrate on the focus of this paper: the comparison of bacterial cell state and bacterial spatial location on treatment outcome. In future papers, the administration of drugs will more closely model the current treatment protocols. In this first iteration of the model, the distribution of antibiotic drug type *i, Drug*_*i*_(**x**, *t*) is governed by a similar equation to that of the oxygen distribution (1), given by

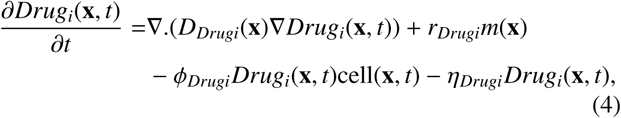

where *D*_*Drugi*_(**x**) is the diffusion coefficient of the drug, *φ*_*Drugi*_ is the uptake rate of the drug, with 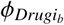 denoting the uptake rate by the bacteria and 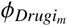 denoting the uptake rate by the infected/chronically infected macrophages. *r*_*Drugi*_ is the drug supply rate by the vascular network and *η*_*Drugi*_ is the drug decay rate. Inside a granuloma structure, the diffusion and supply rate are lower to account for caseum impairing blood vessels and the fact that we know that antibiotic diffusion into granulomata is lower than in normal lung tissue (Kjellsson et al., 2012). Hence the transport properties and delivery rate of the drug are as follows:

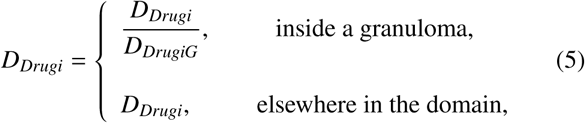

and

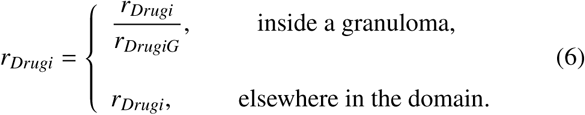

The diffusion rate above is currently based on rifampicin.

To study the efficacy of the drug, we have assumed a threshold drug concentration value, below which the drug has no effect on the bacteria. If the drug reaches a bacterium or infected macrophage when it’s concentration is above this level (which is different for fast- and slow-growing extracellular bacteria and for intracellular bacteria), then the bacterium will be killed and an empty space will be created (this will be described further in section 3.1). Parameters are discussed in section 2.5.

### 2.4 Chemokines

Various molecules are released by macrophages and other immune cells, these molecules act as chemoattractants, attracting other host cells to the site of infection. Although different chemokines perform different roles at various times, for this model, we have chosen to represent the multiple chemokines involved in the immune response as an aggregate chemokine value. Sources of chemokine are derived from infected, chronically infected and activated macrophages (Algood et al., 2003). The distribution of the chemokine molecules, *Ch*(**x**, *t*) is also governed in a similar way to the oxygen and antibiotic:

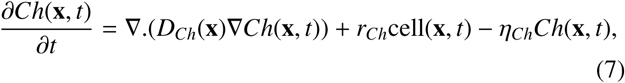

where *D*_*Ch*_(**x**) is the diffusion coefficient of the chemokines, *r*_*Ch*_ is the chemokine supply rate by the macrophages at position **x** at time *t*, with cell(**x**, *t*) = 1 if position **x** is occupied by an infected, chronically infected or activated macrophage at time *t* and zero otherwise. *η*_*Ch*_ is the chemokine decay rate.

### 2.5 Parameter estimation

In order to simulate the model with biologically relevant outcomes, it is important to use accurate parameters values. Most of the parameters are chosen from previous mathematical and experimental papers (see Table 1, Table 2 and Table 3 for a summary of the parameter values).

**Table 1:** Parameter values relating to Equations 2, 3, 5 and 6

|  | Parameter value | Reference |
| --- | --- | --- |
| $D_{OG}$ | 2.7 | (Datta et al., 2015) |
| $r_{OG}$ | 4 | (Datta et al., 2015) |
| $D_{DrugiG}$ | 7.28 (R), 3.85 (H) | (Pienaar et al., 2016) |
| $r_{DrugiG}$ | 4 | (Datta et al., 2015) |

**Table 3:** Parameters. When there is a range for a value, it is set randomly by the model.

| Parameter | Description | Value | Source |
| --- | --- | --- | --- |
| $N$ | Size of grid ($N \times N$) | 100 | (Segovia-Juarez et al., 2004) |
| $N_v$ | Number of blood vessels | 49 | (Cilfone et al., 2013) |
| $\phi_{O_b}$ (Micromoles/cell/hr) | Oxygen consumption rate of bacteria | $20.8 \times 10^{-6}$ | (Sershen et al., 2016) |
| $\phi_{O_{mr}}$ (Micromoles/cell/hr) | Oxygen consumption rate of Mr | $1.15 \times 10^{-7}$ | (Sershen et al., 2016) |
| $\phi_{O_{ma}}$ (Micromoles/cell/hr) | Oxygen consumption rate of Ma | $2.3 \times 10^{-7}$ | (Sershen et al., 2016) |
| $\phi_{O_{mi}}$ (Micromoles/cell/hr) | Oxygen consumption rate of Mi | $3.45 \times 10^{-7}$ | (Sershen et al., 2016) |
| $\phi_{O_{mci}}$ (Micromoles/cell/hr) | Oxygen consumption rate of Mci | $4.6 \times 10^{-7}$ | (Sershen et al., 2016) |
| $\phi_{O_t}$ (Micromoles/cell/hr) | Oxygen consumption rate of T cells | $0.14375 \times 10^{-7}$ | (Sershen et al., 2016) |
| $\phi_{Drug_b}$ (Micromoles/cell/hr) | Antibiotic consumption rate of extracellular bacteria | $2.1 \times 10^{-11}$ | (Pienaar et al., 2015) |
| $\phi_{Drug_{mi}}$ (Micromoles/cell/hr) | Antibiotic consumption rate of infected macrophages | $1.9 \times 10^{-6}$ | (Pienaar et al., 2015) |
| $Rep_f$ (hours) | Replication rate of fast-growing bacteria | 15-32 | (Shorten et al., 2013) |
| $Rep_s$ (hours) | Replication rate of slow-growing bacteria | 48-96 | (Hendon-Dunn et al., 2016) |
| $O_{low}$ (%) | $O_2$ threshold for fast→slow-growing bacteria | 6 | Estimated - see Section 4.1 |
| $O_{high}$ (%) | $O_2$ threshold for slow→fast-growing bacteria | 65 | Estimated - see Section 4.1 |
| $Mr_{init}$ | Initial number of Mr in the domain | 105 | (Cilfone et al., 2013) |
| $MrMa$ | Probability of Mr→Ma (multiplied by no. of T cells in neighbourhood) | 9 | (Cilfone et al., 2013) |
| $N_{ici}$ | Number of bacteria needed for Mi→Mci | 10 | (Cilfone et al., 2013) |
| $N_{cib}$ | Number of bacteria needed for Mci to burst | 20 | (Cilfone et al., 2013) |
| $M_{life}$ (days) | Lifespan of Mr, Mi and Mci | 0-100 | (Van Furth et al., 1973) |
| $Ma_{life}$ (days) | Lifespan of Ma | 10 | (Segovia-Juarez et al., 2004) |
| $t_{moveMr}$ (mins) | Time interval for Mr movement | 20 | (Segovia-Juarez et al., 2004) |
| $t_{moveMa}$ (hours) | Time interval for Ma movement | 7.8 | (Segovia-Juarez et al., 2004) |
| $t_{moveMi}$ (hours) | Time interval for Mi/Mci movement | 24 | (Segovia-Juarez et al., 2004) |
| $Mr_{recr}$ | Probability of Mr recruitment | 0.07 | (Cilfone et al., 2013) |
| $T_{enter}$ | Bacteria needed for T cells to enter the system | 50 | (Cilfone et al., 2013) |
| $T_{recr}$ | Probability of T cell recruitment | 0.02 | (Cilfone et al., 2013) |
| $T_{life}$ (days) | Lifespan of T cells | 0-3 | (Sprent, 1993) |
| $T_{kill}$ | Probability of T cell killing Mi/Mci | 0.75 | (Cilfone et al., 2013) |
| $t_{moveT}$ (mins) | Time interval for T cell movement | 10 | (Segovia-Juarez et al., 2004) |
| $t_{drug}$ (hours) | Time at which drug is administered | 168-2160 | (Asefa and Teshome, 2014; Osei et al., 2015) |
| $DrugKill_f$ (%) | Drug needed to kill fast-growing bacteria | 2 | (Hammond et al., 2015) |
| $DrugKill_s$ (%) | Drug needed to kill slow-growing bacteria | 10 | (Hammond et al., 2015) |
| $DrugKill_{Mi}$ (%) | Drug needed to kill intracellular bacteria | 20 | (Aljayyousi et al., 2017) |

The time step was calculated by considering the fastest process in our system, the oxygen diffusion. The oxygen dynamics are governed by a reaction diffusion equation, where the parameters are chosen from previously published work (Macklin et al., 2012). We assume a oxygen diffusion length scale of *L*=100 *µ*m and a diffusion constant of *D*_*O*_ = 2 × 10^−5^ cm^2^/s (Owen et al., 2004). Using these along with the relation 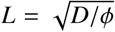, the mean oxygen uptake can be approximately estimated as 0.2 s^-1^. The oxygen supply through the blood vessel is approximately *r*_*O*_ = 8.2 × 10^−3^ mols s^-1^ (Matzavinos et al., 2009).

Supply and diffusion rates inside a granuloma are impaired, as described in Equations 2, 3, 5 and 6. Parameter values are outlined in Table 1.

Nondimensionalisation gives T=0.001 hours and hence each time step is set to Δ*t* = 0.001 hours, with one time step corresponding to 3.6 *s*.

Because oxygen diffusion occurs on a short timescale (of the order of 10*s*), accurately tracking transient variations in oxygen concentration would require a much smaller time step. With *M. tuberculosis* replication rates of between 15 and 96 hours (Shorten et al., 2013; Hendon-Dunn et al., 2016), associated timescales are much faster and hence we assume a quasi-steady profile for oxygen, with the oxygen concentration quickly coming to equilibrium with any changes in the microenvironment.

The simulations are carried out within a two dimensional domain with a grid size *N* = 100, which simulates an area of lung tissue approximately 2 *mm* × 2 *mm*. The step size used is based on the model proposed and developed in (Segovia-Juarez et al., 2004; Marino et al., 2011; Cilfone et al., 2013), with one grid cell corresponding to the approximate diameter of the biggest discrete element in our system, the macrophage (Krombach et al., 1997). The parameters that are used in the equations governing the dynamics of antibiotics and chemokine molecules are chosen in a similar way.

Oxygen is lighter in comparison to the drugs, with a molecular weight of 32 amu (Hlatky and Alpen, 1985), and hence it diffuses faster than most of the drugs and the chemokine molecules. The chemokine molecules diffuse slower than the antibiotics, being heavier than most drugs. One of the drugs under the current study, rifampicin has a molecular weight of822.9 amu (PubChem Compound Database). To obtain or approximate its diffusion coefficient, its molecular mass was compared against the molecular masses of known compounds, as in (Powathil et al., 2012), and consequently taken to be 1.7 × 10^−6^ cm^2^s^-1^. Similar analyses are done with isoniazid, pyrazinamide and ethambutol, with parameter values given in Table 2. The decay rate of these drugs are calculated using the half life values of the drugs obtained from the literature and are also outlined in Table 2.

The threshold drug concentrations, *DrugKill*_*f*_, *DrugKill*_*s*_ and *DrugKill*_*Mi*_, below which the drug has no effect on the TB bacteria have been chosen to be the average density of total drugs delivered through the vessels (total drug delivered/total number of grid points) and the total drug given is kept the same for all drugs types. Values for *DrugKill*_*f*_, *DrugKill*_*s*_ and *DrugKill*_*Mi*_ are based on data arising from *in vitro* experiments and are reported in (Hammond et al., 2015; Aljayyoussi et al., 2017). They are given in Table 3. A relative threshold is chosen here in order to compare the effects of bacterial cell state and location of bacteria, rather than studying any optimisation protocols for drug dosage. As more experimental data becomes available, these thresholds can be refined. Values of 10^−6^ cm^2^s^-1^ to 10^−7^ cm^2^s^-1^ have been reported as diffusion constants for chemokine molecules (Francis and Palsson, 1997). The half-life for IL-8, an important chemokine involved in the immune response of *M. tuberculoisis*, has been shown to be 2-4 hours (Walz et al., 1996). We use a diffusion rate of 10^−6^ cm^2^s^-1^ and a half-life of 2 hours in our simulations.

Other model parameters will be discussed in the next section and are summarised in Table 3.

## 3. Cellular automaton rules

The entire multiscale model is simulated over a prescribed time duration, currently set to 12000 hours (500 days), and a vector containing all grid cell positions is updated at every time step. The oxygen dynamics, chemokine dynamics and drug dynamics are simulated using finite difference schemes.

### 3.1. Rules for the extracellular bacteria

A minimal infectious dose of *M. tuberculosis* has been shown to be of the order of 10 (Capuano et al., 2003). For this reason, we begin our CA simulations with one cluster of 12 bacteria on the grid; 6 fast-growing bacteria and 6 slow-growing bacteria. These initial bacteria replicate following a set of rules and produce a cluster of bacteria on a regular square lattice with no-flux boundary conditions. The fast- and slow-growing bacteria are assigned a replication rate; *Rep*_*f*_ for the fast-growing and *Rep*_*s*_ for the slow-growing. When a bacterium is marked for replication, its neighbourhood of order 3 is checked for an empty space. The neighbourhood type alternates between a Moore neighbourhood and a Von Neumann neighbourhood to avoid square/diamond shaped clusters, respectively. If a space in the neighbourhood exists, a new bacterium is placed randomly in one of the available grid cells. If there are no spaces in the neighbourhood of order 3, the bacterium is marked as ‘resting‘.

At each time step, the neighbourhood of these ‘resting’ bacteria are re-checked so that they can start to replicate again as soon as space becomes available.

As this multiscale model evolves over time, the elements move and interact with each other according to the CA model. The bacteria and host cells also influence the spatial distribution of oxygen since they consume oxygen for their essential metabolic activities. As the bacteria proliferate, the oxygen demand increases creating an imbalance between the supply and demand which will eventually create a state where the bacteria are deprived of oxygen. Bacteria can change between fast-growing and slow-growing states, depending on the oxygen concentration, scaled from 0 to 100, at their location. Fast-growing bacteria where the oxygen concentration is below *O*_*low*_ will become slow-growing, and slow-growing bacteria can turn to fast-growing in areas where the oxygen concentration is above *O*_*high*_ (see Table 3 for these values and section 3.5 for more details).

### 3.2 Rules for the macrophages

There are 4 types of macrophage in our system: resting (Mr), active (Ma), infected (Mi) and chronically infected (Mci). There are *Mr*_*init*_ resting macrophages randomly placed on the grid at the start of the simulation. These resting macrophages can become active when T cells are in their Moore neighbourhood, with probability *MrMa* multiplied by the number of T cells in the neighbourhood. When active macrophages encounter extracellular bacteria, they kill the bacteria. If the resting macrophages encounter bacteria, they become infected and can become chronically infected when they phagocytose more than *N*_*ici*_ bacteria. Chronically infected macrophages can only contain *N*_*cib*_ intracellular bacteria, after which they burst. Bursting macrophages distribute bacteria randomly into their Moore neighbourhood of order 3 and the grid cell where the macrophage was located becomes caseum.

While the oxygen and antibiotics enter the system via the blood vessel network, the chemokines are secreted by the infected, chronically infected and activated macrophages. Macrophages move in biased random walks, with probabilities calculated as a function of the chemokine concentration of its Moore neighbourhood. Resting, infected and chronically infected macrophages are randomly assigned a lifespan, *M*_*li* *f* *e*_ days, and active macrophages live for *Ma*_*li* *f* *e*_ days. Resting macrophages move every *t*_*moveMr*_ minutes, active macrophages move every *t*_*moveMa*_ hours and infected/chronically infected macrophages move every *t*_*moveMi*_ hours. Resting macrophages are recruited from the blood vessels with a probability of *Mr*_*recr*_.

### 3.3 Rules for the T cells

The T cells enter the system once the extracellular bacterial load reaches *T*_*enter*_ and move in a biased random walk, similar to the macrophages. T cells are recruited from the blood vessels with a probability *T*_*recr*_. They live for *T*_*li* *f* *e*_, and move every *t*_*moveT*_ minutes. Activated T cells are immune effector cells that can kill chronically infected macrophages. If a T cell encounters an infected or chronically infected macrophage, it kills the macrophage (and all intracellular bacteria) with probability *T*_*kill*_ and that grid cell becomes caseum. T cells also activate resting macrophages when they are in their Moore neighbourhood, with probability *MrMa* multiplied by the number of T cells in the neighbourhood.

### 3.4 Rules for the Antibiotics

If the immune process does not clear the infection, drugs are administered at *t*_*drug*_ hours, a randomly chosen time between two values (see Table 3). This mimics the variability in time that patients seek medical attention for their disease (Asefa and Teshome, 2014; Osei et al., 2015). The antibiotics are administered for a period of 4380 hours (six months). The antibiotic kills the bacteria when the concentration is over *DrugKill*_*f*_ or *DrugKill*_*s*_, for the fastand slow-growing bacteria respectively. The antibiotics can also kill intracellular bacteria, by killing the bacteria contained within the infected/chronically infected macrophages if the concentration is over *DrugKill*_*Mi*_. These macrophages then return to resting macrophages.

### 3.5. Oxygen thresholds for bacterial cell states

In order to choose values for *O*_*low*_ and *O*_*high*_, we conducted some test simulations for 120 hours, where no macrophages were present. This allowed us to alter both the oxygen switching thresholds and see the effect it had on the bacteria. 90 simulations were run in total, with a range of both parameters being tested. Table 4 describes the outcome of these simulations:

**Table 4:** Oxygen switch

| | $O_{low} = 3$ | $O_{low} = 6$ | $O_{low} = 9$ |
| --- | --- | --- | --- |
| $O_{high} = 55$ | All slow-growing bacteria switched to fast-growing in all 10 simulations by $t = 10$ hours | In 9 of the 10 simulations, all slow-growing bacteria changed to fast-growing by $t = 10$ hours | In all 10 simulations, bacteria was all the same cell state by $t = 10$ hours: in 4 simulations all were slow-growing, in 6 simulations all were fast-growing |
| $O_{high} = 65$ | No bacteria change state in any of the 10 simulations | In 7 of the simulations, some bacteria changed state: 4 from fast-growing to slow-growing and 3 from slow-growing to fast-growing (mean time for first switch was $t=21$ hours). In 3 of the simulations, no bacteria changed state. | All fast-growing bacteria become slow-growing by $t = 10$ hours in all 10 simulations |
| $O_{high} = 75$ | No bacteria change state in any of the 10 simulations | No bacteria changed state in any of the 10 simulations | All fast-growing bacteria become slow-growing by $t = 10$ hours in all 10 simulations |

Figure 5 shows representative examples of these test simulations, where fast- and slow-growing bacteria are shown (by the blue/cyan lines, respectively), with *O*_*high*_ fixed at 65 and *O*_*low*_ = 3, 6, 9. In (a) we see the effect of having a lower threshold for fast-growing bacteria to become slow-growing, with *O*_*low*_ = 3, in (b) *O*_*low*_ = 6 and in (c), *O*_*low*_ has a higher threshold of 9. In simulation (a), the bacteria do not change state during the entire simulation, Figure 5 (a) supports this as we do not see an increase in the cyan line that corresponds with a drop in the blue. Simulation (b) has *O*_*low*_ = 6 and here we see some transfer from fast- to slow-growing during the 120 hours. For simulation (c), however, the fast-growing bacteria transfer to slow-growing almost immediately.

**Figure 5:**
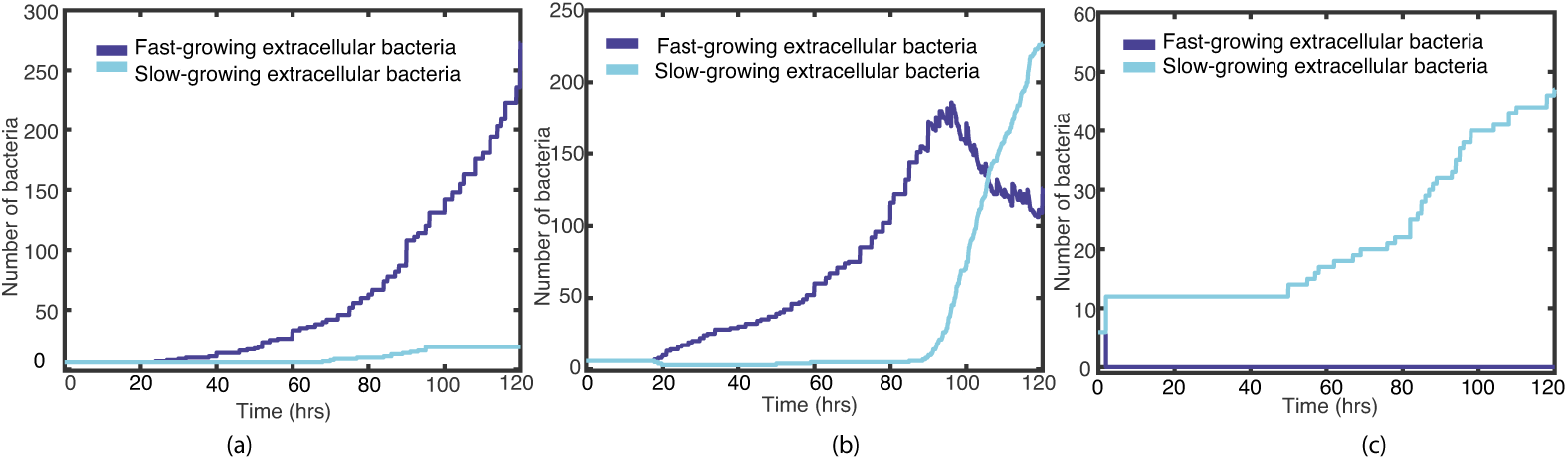
Plots of the fast-(blue) and slow-growing (cyan) extracellular bacteria for the first 120 hours, with *O*_*high*_ fixed at 65 and (a) *O*_*low*_ = 3, (b) *O*_*low*_ = 6 and (c) *O*_*low*_ = 9.

Similarly, Figure 6 shows representative examples from varying *O*_*high*_, where *O*_*low*_ is held at 6 and *O*_*high*_ = 55, 65, 75. In simulation (a), where *O*_*high*_ = 55, the slow-growing bacteria all change to fast-growing very near to the beginning of the simulation. Simulation (b) shows some bacteria changing state when *O*_*high*_ = 65, and simulation (c) shows no transfer from fastto slow-growing when *O*_*high*_ = 75.

**Figure 6:**
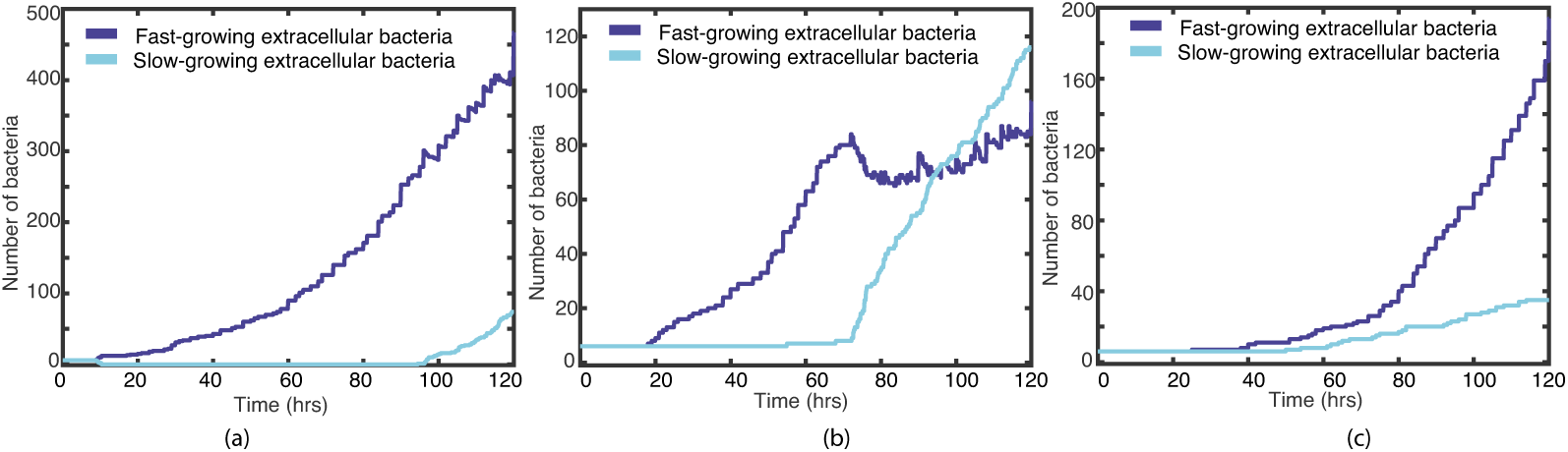
Plots of the fast-(blue) and slow-growing (cyan) extracellular bacteria for the first 120 hours, with *O*_*low*_ fixed at 6 and (a) *O*_*high*_ = 55, (b) *O*_*high*_ = 65 and (c) *O*_*high*_ = 75.

These test simulations support us choosing *O*_*low*_ = 6 and *O*_*high*_ = 65.

## 4. Simulation results

In order to study the relative importance of bacterial cell state and initial spatial location of bacteria, we study two scenarios: one with a fixed, uniform blood vessel distribution and initial bacterial locations (see four examples in Figure 7 (a)), and another where the vessel distribution and the initial locations of the extracellular bacteria are determined randomly for each simulation (see examples in Figures 9-11 (a)). We run a total of 120 simulations for 500 days: 20 simulations for the ‘fixed’ scenario and 100 simulations for the ‘random’ scenario. These simulations were run on servers that have dual Intel Xeon E5-2640 CPUs and 128GB RAM. Each simulation took approximately 8 hours to run.

**Figure 7:**
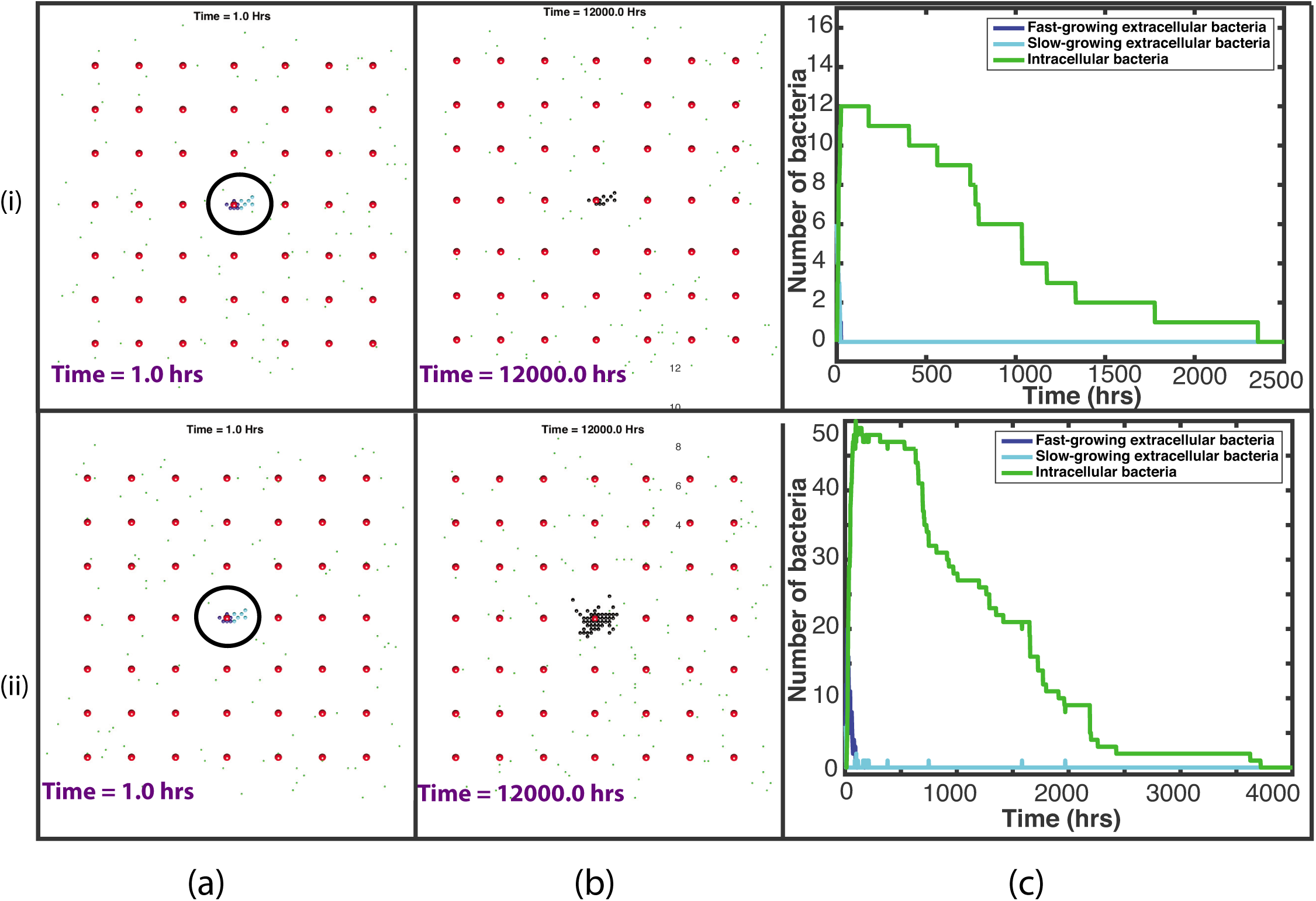
Plots showing the outcome of two representative simulations, (i)-(ii), with the fixed vessel distribution and initial bacterial location. (a) and (b) are plots of the spatial distribution of all elements: at the start of the simulation, with the initial bacterial cluster highlighted by the black circle (a) and at the end of the simulation (b). Red circles depict the blood vessels, black circles depict the caseum, blue circles show the fast-growing extracellular bacteria, cyan circles show the slow-growing extracellular bacteria, green dots depict macrophages (with darker green for the infected/chronically infected macrophages) and yellow dots depict the T cells. Plots of bacterial numbers are shown in (c), depicting fast-growing extracellular bacteria (dark blue), slow-growing extracellular bacteria (cyan) and intracellular bacteria (green). Note that individual figures are available in Supplementary material for closer inspection.

**Figure 9:**
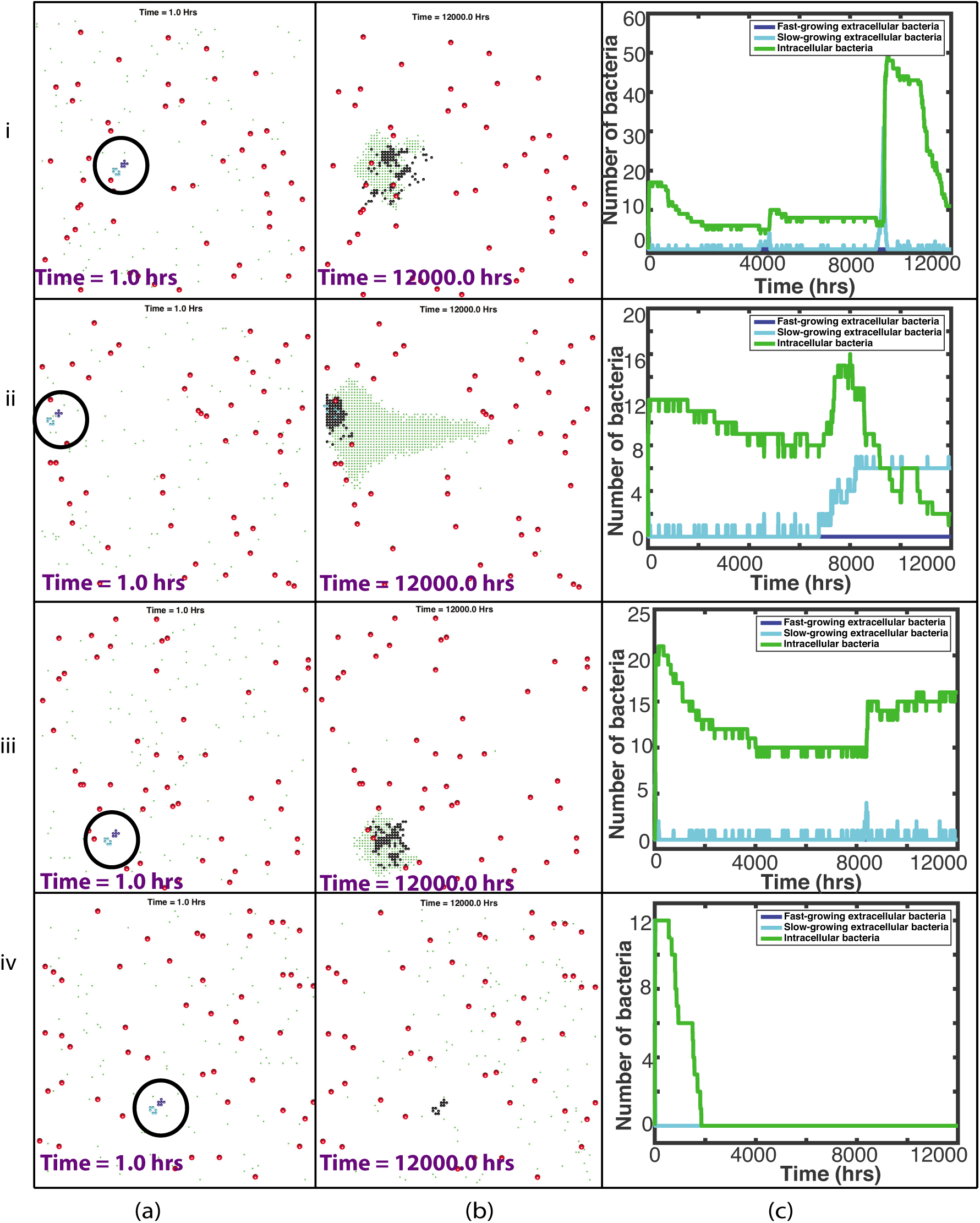
Plots showing the outcome of four representative simulations, (i)-(iv), of contained simulations, with a randomly placed vessel distribution and initial bacterial location. (a)-(b) are plots of the spatial distribution of all elements: at the start of the simulation, with the initial bacterial cluster highlighted by the black circle (a) and at the end of the simulation (b). Red circles depict the blood vessels, black circles depict the caseum, blue circles show the fast-growing extracellular bacteria, cyan circles show the slow-growing extracellular bacteria, green dots depict macrophages (with darker green for the infected/chronically infected macrophages) and yellow dots depict the T cells. Plots of bacterial numbers are shown in (d), depicting fast-growing extracellular bacteria (dark blue), slowgrowing extracellular bacteria (cyan) and intracellular bacteria (green). Note that individual figures are available in Supplementary material for closer inspection.

### 4.1 Fixed blood vessels and initial bacterial cluster

20 simulations were run with the same initial distribution of *N*_*v*_ blood vessels and with one bacterial cluster of 6 fast-growing bacteria and 6 slow-growing bacteria, located in the centre of the grid. Figure 7 shows this initial set up. In this ‘fixed’ spatial configuration, there was 1 blood vessel located within a 0.1 mm radius of the bacterial cluster, situated 1 grid cell away (0.02 mm). All 20 simulations resulted in containment of the disease (which we define as fewer than 10 extracellular bacteria at the end of the 500 days), with the macrophages and T cells preventing disease progression. In all 20 simulations no bacteria remained at the end of the 500 days. As a granuloma develops in the simulations, caseous cells are created at the centre. At 500 days, the simulations had a range of 11-49 caseous grid cells with a median value of 13.5. In only one simulation the total bacterial load exceeded *T*_*enter*_ and therefore T cells only appeared in one simulation. See Table 1 in supplementary material for summary statistics for these 20 simulations. Supplementary material also contains a document with plots showing the dynamics of the T cells.

As can be seen in Figure 7(i)(b), at the end of this fixed simulation, there are 10 caseous grid cells remaining, with all bacteria eradicated. In the line plot, Figure 7(i)(c), we see that the macrophages effectively phagocytose the bacteria early on in the simulation and the infected macrophages gradually die out over 2500 hours (around 100 days). In the model, as a macrophage reaches the end of its lifespan, the grid cell becomes caseum and the intracellular bacteria contained within is able to burst out and grow extracellularly. This can only happen, however, if there is space for the bacterium to move to. If instead, as in this case, the immune response is effective in controlling the disease with resting macrophages surrounding the infection, there is no space for these intracellular bacteria to re-emerge as extracellular bacteria, and the bacterium dies.

Figure 7(ii) depicts the outcome of another example simulation with a starting ‘fixed’ distribution. Figure 7(ii)(b) shows the end of this simulation with 49 caseous grid cells. If we look at the line plot in Figure 7(ii)(c), we can see a more eventful simulation. The bacteria start to grow at the beginning of the simulation and we see a more gradual immune containment of the infection. This is the one simulation where T cells entered the system to assist in this containment. Because of this, the T cells are also responsible for killing infected macrophages and they also activate the macrophages, which also contribute to the killing of the infected macrophages. In Figure 7(ii)(c) we also see incidences of intracellular bacteria successfully moving out of the macrophages as they die naturally. These can be seen as small spikes in the slow-growing line. In many of these cases, however, these newly escaped extracellular bacteria are quickly phagocytosed again. Eventually, by around 3700 hours (around 150 days), this infection is completely eradicated.

Figure 8 shows summary plots of the 20 fixed simulations, with the mean bacterial numbers shown by the solid lines and 95% confidence intervals shown by the dashed lines. Figure 8(i) shows the entire 12000 hours and Figure 8(ii) shows only until 200 hours, as this is where most activity takes place. We can see from these plots that there is not a great amount of variation between the 20 simulations, with small differences in bacterial numbers. It is clear also that most of the variability takes place close to the start of the simulations. This confirms our conclusions that with a fixed distribution of blood vessels and initial bacterial placement, there is very little difference in the outcome by the end of the simulations.

**Figure 8:**
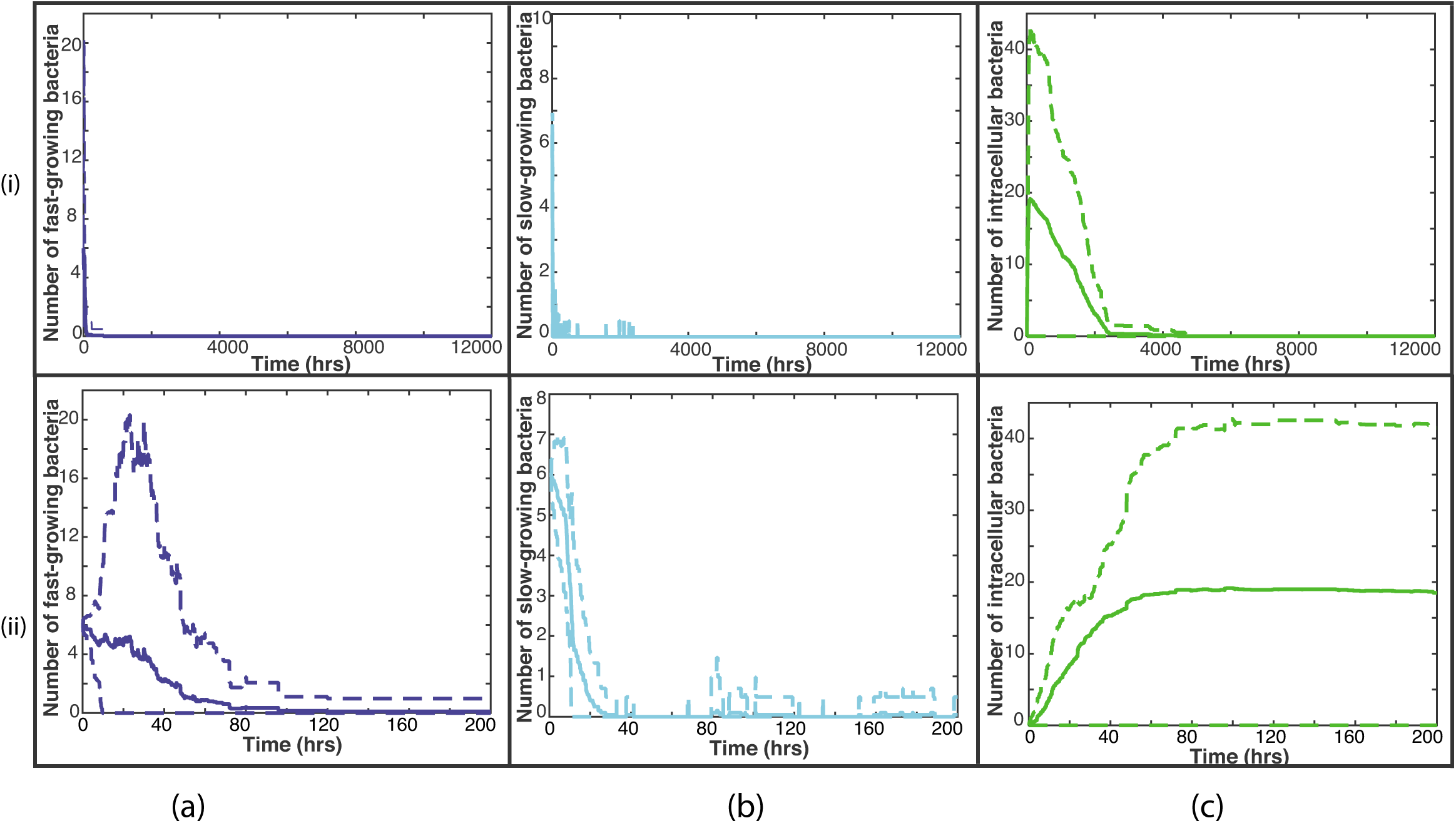
Summary plots of all 20 fixed simulations. The solid line represents the mean bacterial numbers and the dashed lines show the 95% confidence intervals. (i) shows the bacterial numbers for the entire 12000 hours, whereas (ii) shows until time 200 hours, where most activity takes place. The fast-growing bacteria are shown by the blue lines in (a), the slow-growing bacteria are shown by the cyan lines in (b) and the intracellular bacteria by the green lines in (c).

### 4.2. Random blood vessels and initial bacterial cluster

100 simulations were run with a random distribution of blood vessels and a random location for a bacterial cluster. The bacterial cluster consisted of 6 fast-growing bacteria and 6 slow-growing bacteria, as in the fixed scenario. 90 simulations resulted in containment of the disease. Again we define containment as fewer than 10 extracellular bacteria at the end of the 500 days. Ten simulations had a number of slow-growing bacteria remaining at 500 days, with a range 1-6. All of these remaining extracellular bacteria were surrounded by caseous grid cells. The other 80 simulations had no extracellular bacteria remaining. 29 out of the 90 simulations had intercellular bacteria, with a range 1-41. Of these 90 ‘contained’ simulations, 32 still had a small number of either extracellular or intracellular bacteria remaining at the end of the simulation (but fewer than ten extracellular bacteria) and hence these would be termed ‘latently infected‘, where the disease is capable of progressing at a later stage. Figure 9 shows four representative examples of these 90 simulations that were contained.

In Figure 9 (i)(b), we see an example of a simulation where the infection has been contained, with no extracellular bacteria and only 11 intracellular bacteria remaining. There is also a granuloma visible at 500 days, containing 102 caseous grid cells. The line plot shown in Figure 9 (i) (c) describes how the immune system has contained this infection. In the 500 days of the simulation, we see numerous spikes in the extracellular bacteria, as infected macrophages die and the intracellular bacteria are released. In most circumstances, this re-emergent infection is quickly controlled. At roughly 9000 hours, however, the slow-growing bacteria begin to grow again. It is at this point in the simulation that T cells are recruited, which helps to regain control of the infection.

Figure 9 (ii) shows another example of a contained infection, where no treatment was needed. In this simulation, there are 6 extracellular bacteria remaining at the end of the 500 days. However, as these are surrounded by caseous material within the granuloma structure, the infection is controlled and the bacteria cannot grow.

In Figure 9 (iii), we see a similar picture to that of shown in Figure 9 (i). The difference here is that there was no T cell recruitment needed to control the infection. By the end of the simulation, we see in Figure 9 (iii)(c) that there are 16 intracellular bacteria remaining.

Figure 9 (iv) depicts a situation where the infection is controlled efficiently by the host immunity. Figure 9 (iv)(b) shows no bacteria remaining with only 12 caseous grid cells.

In 10 simulations, the immune response was not able to contain the disease within the granuloma and active disease developed. Antibiotics were then administered at *t*_*drug*_ hours. In seven of these simulations, the treatment was ‘successful‘, where we define success as fewer than ten extracellular bacteria at the end of 500 days. Four of these simulations had between 1 and 8 slow-growing extracellular bacteria remaining at 500 days. These extracellular bacteria were all surrounded by caseous grid cells. Although these four simulations are deemed ‘successful‘, as there are a small number of bacteria remaining, these cases are capable of relapsing. In the seven successful simulations, the majority of the bacteria are killed by the antibiotics (mean value of 75.1%). Four representative examples of these ‘successfully treated cases’ are shown in Figure 10.

**Figure 10:**
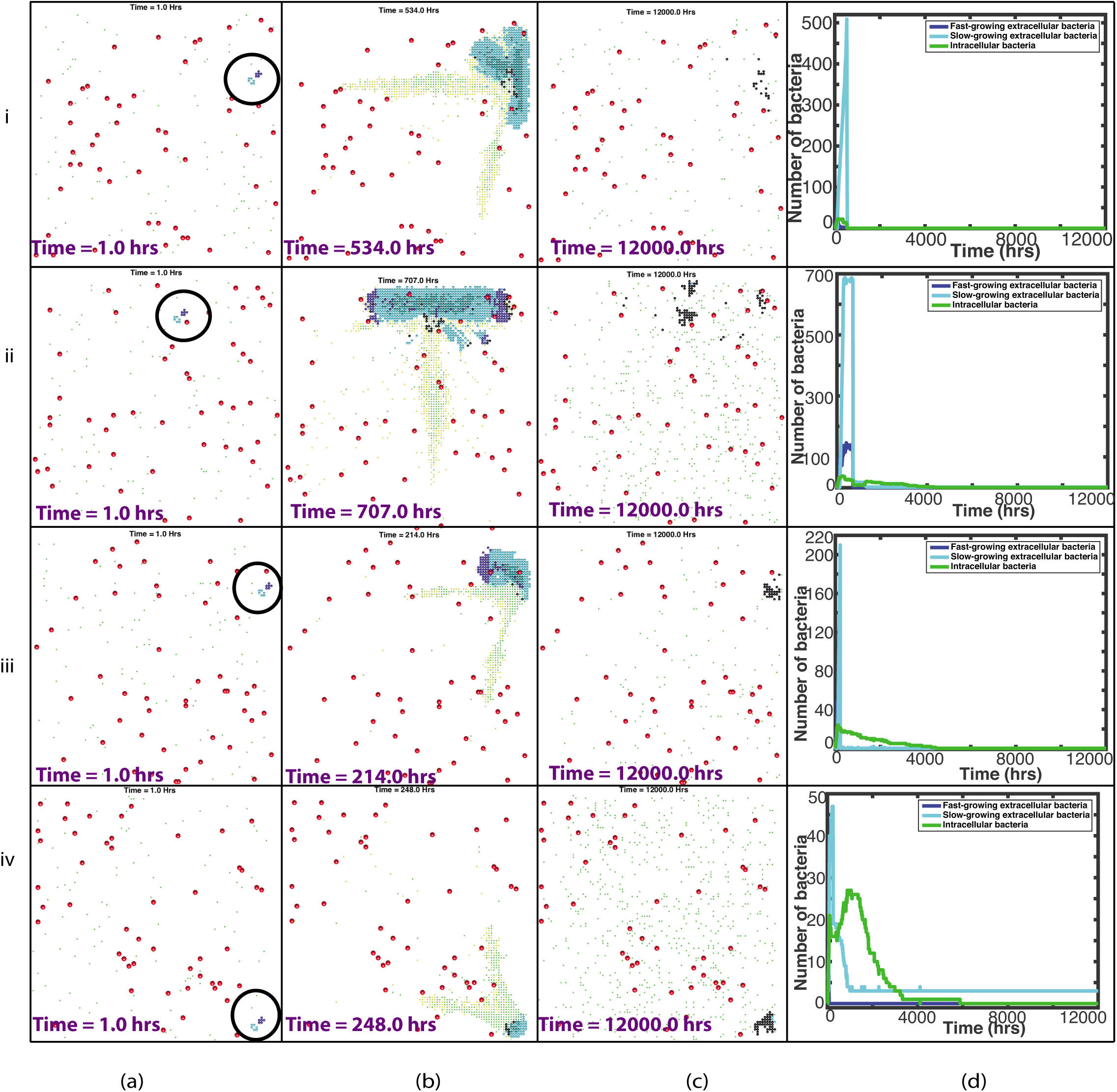
Plots showing the outcome of four representative simulations, (i)-(iv), which were ‘successfully treated‘, with a randomly placed vessel distribution and initial bacterial location. (a)-(c) are plots of the spatial distribution of all elements: at the start of the simulation, with the initial bacterial cluster highlighted by the black circle (a), just before the drug enters the system (b) and at the end of the simulation (c). Red circles depict the blood vessels, black circles depict the caseum, blue circles show the fast-growing extracellular bacteria, cyan circles show the slow-growing extracellular bacteria, green dots depict macrophages (with darker green for the infected/chronically infected macrophages) and yellow dots depict the T cells. Plots of bacterial numbers are shown in (d), depicting fast-growing extracellular bacteria (dark blue), slow-growing extracellular bacteria (cyan) and intracellular bacteria (green). Note that individual figures are available in Supplementary material for closer inspection.

Figure 10 (i) shows an example of a successfully treated simulation. Both types of extracellular bacteria grow rapidly at the start of the simulation, with the immune cells unable to control the infection. As the bacteria grow, they start to consume more and more oxygen, reducing the availability and causing the fast-growing bacteria to switch state. We see the clear effects of the antibiotics as they enter the system at 535 hours, with 85.8% of the total bacteria in the system being killed by the antibiotics at this time. No bacteria remain at the end of this simulation.

Figure 10 (ii) shows another example of a successfully treated simulation. Again, we see the sharp reduction of bacterial load as the antibiotics enter the system at 708 hours, with 80.6% of the total bacteria in the system being killed by the antibiotics. Shortly after this, the host immunity controls the remaining infection. Only 1 slow-growing bacteria remains at the end of the simulation, surrounded by casous grid cells.

In Figure 10 (iii), antibiotics were administered at 215 hours, which we see in Figure 10 (iii)(d) controls the infection, killing 88.9% of the total bacteria.

Figure 10 (iv) shows an example where, although the antibiotics were used and were effective, the immune response was actually responsible for the majority of the killing (63.5%). This is in part due to that fact that antibiotics were given early, at 249 hours when the bacterial burden was not particularly high. The remaining three simulations, were deemed ‘unsuccessful‘. In these simulations there were 16, 241 and 354 slow-growing bacteria remaining at 500 days. The latter two simulations are shown in Figure 11.

**Figure 11:**
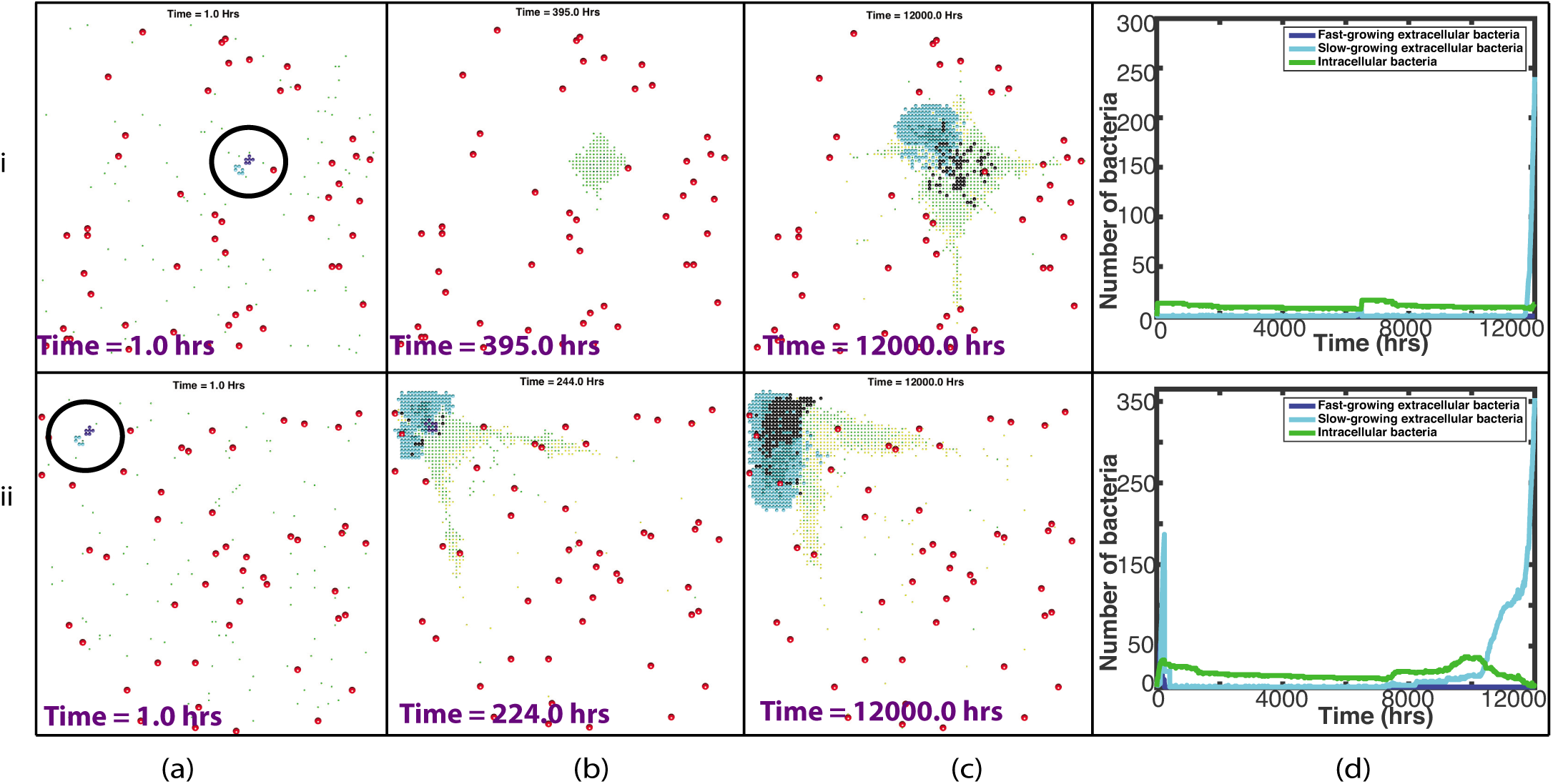
Plots showing the outcome of two representative simulations, (i)-(ii), which were ‘unsuccessfully treated‘, with a randomly placed vessel distribution and initial bacterial location. (a)-(c) are plots of the spatial distribution of all elements: at the start of the simulation, with the initial bacterial cluster highlighted by the black circle (a), just before the drug enters the system (b) and at the end of the simulation (c). Red circles depict the blood vessels, black circles depict the caseum, blue circles show the fast-growing extracellular bacteria, cyan circles show the slow-growing extracellular bacteria, green dots depict macrophages (with darker green for the infected/chronically infected macrophages) and yellow dots depict the T cells. Plots of bacterial numbers are shown in (d), depicting fast-growing extracellular bacteria (dark blue), slow-growing extracellular bacteria (cyan) and intracellular bacteria (green). Note that individual figures are available in Supplementary material for closer inspection.

Figure 11 (i) shows a case where the infection was controlled by the host immunity for almost the entire simulation. Near the end of the 500 days, however, the intracellular bacteria escape as the macrophages reach the end of their lives and these newly extracellular bacteria grow. This is an example of a ‘latent’ case of tuberculosis where the infection reactivates at a later date.

Finally, Figure 11 (ii) shows an example of a simulation where treatment was received and was initially ‘successful’ but, as it can be seen in Figure 11 (ii)(d), at around 8000 hours, extracellular bacteria begin to grow as dying macrophages release their intracellular bacteria. The slow-growing bacteria then continue to grow until the end of the simulation. This is an example of a relapse.

Figure 12 shows summary plots of the 100 random simulations, with the mean bacterial numbers shown by the solid lines and 95% confidence intervals shown by the dashed lines. Figure 12(i) shows the entire 12000 hours and Figure 12(ii) shows only until 200 hours, as this is where most activity takes place. In contrast to the fixed summary plots, we see much more variation in bacterial numbers, especially for the slow-growing bacteria. We also note that this variability remains for the entire 12000 hours. This further highlights the increased variability in outcome when we investigate a random placement of blood vessels and bacterial cluster.

**Figure 12:**
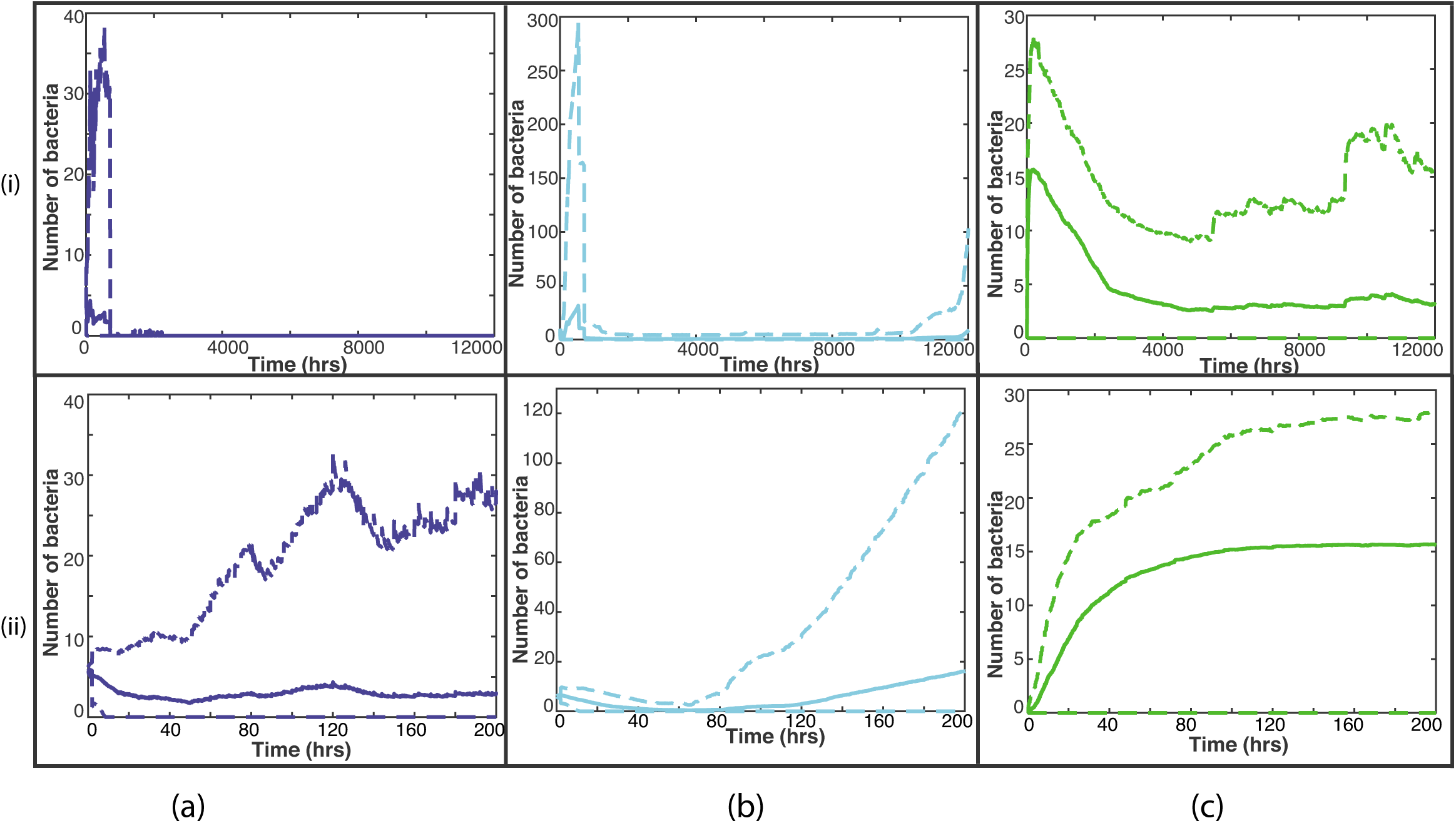
Summary plots of all 100 random simulations. The solid line represents the mean bacterial numbers and the dashed lines show the 95% confidence intervals. (i) shows the bacterial numbers for the entire 12000 hours, whereas (ii) shows until time 200 hours, where most activity takes place. The fast-growing bacteria are shown by the blue lines in (a), the slow-growing bacteria are shown by the cyan lines in (b) and the intracellular bacteria by the green lines in (c).

Table 2 in supplementary material shows summary statistics for these 100 random simulations. Supplementary material also contains a document with plots of the dynamics of the T cells.

In the 90 ‘contained’ simulations, the median distance of initial bacterial cluster to nearest blood vessel source is 0.1 mm, with a median value of 1 blood vessel source within a 0.1 mm radius of the bacteria. In contrast, the other 10 simulations which were not contained by host immunity, the median distance to the closest blood vessel source is 0.16 mm, with a median value of 0 blood vessel sources within a 0.1 mm radius of the bacteria. Comparing the contained cases with the treated cases, a 1-tailed student t-test gives a p value of *p* = 0.014 for proximity to closest blood vessel and *p* = 4.6 × 10^−5^ for number of blood vessels within a 0.1 mm radius. Hence we have shown that these groups are statistically different at the 5% level. This seems to suggest that, if the initial bacteria are located further away from the blood vessels, the less likely it is that the host immune response will contain the infection.

## 5. Discussion

Individual-based models have already been shown to be useful in understanding tuberculosis disease progression (Segovia-Juarez et al., 2004; Marino et al., 2011; Cilfone et al., 2013; Pienaar et al., 2015, 2016; Sershen et al., 2016). Here we have built a hybrid cellular automaton model that incorporates oxygen dynamics, which allows bacteria to change state, and includes antibiotic treatments. In addition to focusing on bacterial cell state, we also investigate changes in spatial location of the bacteria and their influences on disease outcome.

We have shown that position of bacteria in relation to the source of drugs alters the outcome of simulations. When analysing the 20 simulations with a fixed, uniform blood vessel distribution, we see that there is very little difference between the simulations, with the host immunity containing the infection in all 20 cases. For the random distribution of blood vessels, the location of the bacterial cluster was an important factor in determining disease outcome. In the simulations where treatment was necessary, the initial bacteria were usually further away from the blood vessel sources than those that were contained by the immune response. The blood vessels act as sources for the immune cells and so when the bacteria are located further away from these sources, it can take a longer time for the immune cells to respond to and attempt to control the infection. This extra time can give the bacteria time to grow, making it harder for the host immunity to contain the disease, thus in the majority of cases, antibiotics are required to reduce the bacterial burden. We also note that in many of the simulations where the initial bacterial cluster is located near the edge of the computational domain (examples shown in Figure 10), the host immune response tends to find it harder to contain the infection. This is because the host cells cannot surround the bacteria and therefore a complete granuloma cannot develop. Although this is a flaw in the model design, it does illustrate the importance of effective granuloma formation. In future iterations of the model we could consider a larger domain or look at alternative boundary conditions to compare the effect.

In the 100 simulations that used a random distribution of blood vessels and bacteria, we found that 90 (90%) of them were contained by the host immunity but 32 of these (36%) still had some bacteria remaining, and are therefore latently infected, their disease capable of reactivating at a later date. 10 of the 100 simulations (10%) required treatment and 7 of these (70%) had favourable outcomes, with fewer than ten extracellular bacteria remaining at the end of 500 days. The remaining three simulations had more than ten extracellular bacteria remaining at the end of the simulation. Two of these cases were treated with antibiotics, which reduced the bacterial load dramatically. In one case, only 16 bacteria were remaining and these bacteria were situated within a granuloma. The other case depicts a relapse, where treatment was initially successful but infection started to grow again post treatment. The last case is an example of a latently infected individual whose disease reactivates just before the end of the simulation. These percentages are comparable with those described in Figure 1 and from epidemiological data from the WHO on the rates of latent TB disease (World Health Organisation, 2013), with 90% of the cases resulting in containment of the disease but with many capable of later relapse. We also show a case where an early clearance of the bacterial load still ended up with relapse of the disease, this is consistent with observations from the REMoxTB trial, reported in (Phillips et al., 2016). Our model simulations also show that in the cases where bacteria remain at the end of the simulation, they are slow-growing. This finding of persistent bacteria is also consistent with the literature describing persistent TB infection in *in vivo* models (Hu et al., 2000; Manina et al., 2015).

An important feature of our model is that of caseation: when infected macrophages burst or die, or T cells kill infected macrophages, that grid cell becomes caseum. Hence, as macrophages move chemotactically towards the clusters of bacteria, a caseous granuloma starts to form and this caseum inhibits drug diffusion. We have shown that in simulations where bacteria are surrounded by caseum, they often remain at the end of the simulation. This emphasises the importance of caseous necrosis on the outcome of therapy. The implications of caseum have already been demonstrated (Grosset, 1980) and our simulations confirm the importance of this type of necrotic breakdown. Other hybrid models, such as (Pienaar et al., 2015), have also included caseum, with non-replicating bacteria residing in the necrotic tissue and also call attention to this important feature in terms of treatment outcome. With recent experimental studies such as (Sarathy et al., 2018), we now have the available data to explore the drug susceptibility of bacteria in caseum.

We have also shown that bacterial cell state has an impact on simulations, which is a characteristic that is only just starting to be understood. All simulations that have extracellular bacteria remaining at the end of 500 days are slow-growing bacteria. This is in part because any treatment received tends to kill off the faster-growing bacteria more quickly, perhaps leaving behind a slower-growing population. These bacteria are also often located inside a granuloma, where the oxygen supply and diffusion rate is impaired, which favours slower-growing phenotypes. There are relatively few publications that define the susceptibility of slow-growing mycobacteria in relation to the standard or new anti-tuberculosis drugs and particular emphasis should be placed on such antibiotics that can penetrate well into lesions.

Our preliminary simulations also highlight the importance of spatial location of the bacteria. Perhaps it is thought obvious that spatial location of the bacteria is a key factor in treatment outcome but previous mathematical models to date have not identified this fact. Studies have focused on pharmacokinetic (PK) based on serum and simulations of epithelial lining fluid (ELF) bronchoalveolar lavage. Our modelling has shown that anatomical considerations are important when chronic infection creates an anaerobic environment and fibrosis around cavities. Treatment is compounded further by bacterial cell state, which increases functional MIC of bacteria that can be more difficult to kill due to poor penetration.

Future models could address this with enhanced understanding of the effect of dormancy or phenotypic resistance. This indicates the importance of work to define lesional PK (Prideaux et al., 2015; Via et al., 2015). In future iterations of the model will also include more anatomical and immunological complexity, for example, airways will be added to the domain to explore its effects and more than one T cell will be integrated into the model. We could also explore the effect of fibrosis and cavity formation on outcome, building on recent concepts on lesional drug concentrations (Prideaux et al., 2015; Via et al., 2015). In addition, we will model liquefaction, which will be important to allow the release of ‘trapped’ bacteria, thus allowing us to further investigate relapse cases. As our understanding of *M. tuberculosis* cell state increases we will also be able to refine our parameter estimates of this characteristic and build a better model. In future models we will also investigate the effect of allowing more than one element to occupy each grid cell. In this model the simple drug diffusion that is modelled is sufficient for our initial investigations. In future models we will integrate a PK/PD treatment model for combination therapy so that we can investigate the relative importance of front-line drugs and their role in targeting the various bacteria.

Sputum culture conversion during treatment for tuberculosis has a limited role in predicting the outcome of treatment for individual patients (Phillips et al., 2016), so spatial models that explore TB infection and treatment in the lung are needed if we are to increase our understanding of patient outcome. In this work we have shown, using an individual-based model, that a spatial model allows us to explore many unanswered questions in TB.

## Acknowledgements

This work was supported by the Medical Research Council [grant number MR/P014704/1] and the PreDiCT-TB consortium (IMI Joint undertaking grant agreement number 115337, resources of which are composed of financial contribution from the European Union’s Seventh Framework Programme (FP7/2007-2013) and EFPIA companies’ in kind contribution.

## References

Ahmad, S., 2010. Pathogenesis, immunology, and diagnosis of latent mycobacterium tuberculosis infection. Clinical and Developmental Immunology 2011.

Alarcón, T., Byrne, H. M., Maini, P. K., 2003. A cellular automaton model for tumour growth in inhomogeneous environment. J. Theor. Biol. 225 (2), 257–274.

Algood, H. M. S., Chan, J., Flynn, J. L., 2003. Chemokines and tuberculosis. Cytokine Growth Factor Rev. 14 (6), 467–477.

Aljayyoussi, G., Jenkins, V. A., Sharma, R., Ardrey, A., Donnellan, S., Ward, S. A., Biagini, G. A., 2017. Pharmacokinetic-pharmacodynamic modelling of intracellular mycobacterium tuberculosis growth and kill rates is predictive of clinical treatment duration. Scientific Reports 7 (1), 502.

Asefa, A., Teshome, W., 2014. Total delay in treatment among smear positive pulmonary tuberculosis patients in five primary health centers, southern ethiopia: a cross sectional study. PloS one 9 (7), e102884.

Baron, V. O., Chen, M., Clark, S. O., Williams, A., Hammond, R. J., Dholakia, K., Gillespie, S. H., 2017. Label-free optical vibrational spectroscopy to detect the metabolic state of m. tuberculosis cells at the site of disease. Scientific Reports 7 (1), 9844.

Canetti, G., et al., 1955. The tubercle bacillus in the pulmonary lesion of man. The´ Tubercle Bacillus in the Pulmonary Lesion of Man.

Capuano, S. V., Croix, D. A., Pawar, S., Zinovik, A., Myers, A., Lin, P. L., Bissel, S., Fuhrman, C., Klein, E., Flynn, J. L., 2003. Experimental mycobacterium tuberculosis infection of cynomolgus macaques closely resembles the various manifestations of human m. tuberculosis infection. Infection and immunity 71 (10), 5831–5844.

Cilfone, N. A., Perry, C. R., Kirschner, D. E., Linderman, J. J., 2013. Multi-scale modeling predicts a balance of tumor necrosis factor-α and interleukin-10 controls the granuloma environment during mycobacterium tuberculosis infection. PLoS One 8 (7), e68680.

Das¸u, A., Toma-Das¸u, I., Karlsson, M., 2003. Theoretical simulation of tumour oxygenation and results from acute and chronic hypoxia. Phys. Med. Biol. 48 (17), 2829.

Datta, M., Via, L. E., Chen, W., Baish, J. W., Xu, L., Barry 3rd, C. E., Jain, R. K., 2015. Mathematical model of oxygen transport in tuberculosis granulomas. Ann. of Biomed. Eng., 1–10.

Dormann, S., Deutsch, A., 2002. Modeling of self-organized avascular tumor growth with a hybrid cellular automaton. In Silico Biol. 2 (3), 393–406.

Fox, W., 1981. Whither short-course chemotherapy? British journal of diseases of the chest 75 (4), 331–357.

Fox, W., Ellard, G. A., Mitchison, D. A., 1999. Studies on the treatment of tuberculosis undertaken by the british medical research council tuberculosis units, 1946–1986, with relevant subsequent publications. The International Journal of Tuberculosis and Lung Disease 3 (10), S231–S279.

Francis, K., Palsson, B. O., 1997. Effective intercellular communication distances are determined by the relative time constants for cyto/chemokine secretion and diffusion. Proc. Natl. Acad. Sci. 94 (23), 12258–12262.

Gerlee, P., Anderson, A. R., 2007. An evolutionary hybrid cellular automaton model of solid tumour growth. J. Theor. Biol. 246 (4), 583–603.

Gillespie, S. H., Crook, A. M., McHugh, T. D., Mendel, C. M., Meredith, S. K., Murray, S. R., Pappas, F., Phillips, P. P., Nunn, A. J., 2014. Four-month moxifloxacin-based regimens for drug-sensitive tuberculosis. N. Engl. J. Med 371 (17), 1577–1587.

Grosset, J., 1980. Bacteriologic basis of short-course chemotherapy for tuberculosis. Clin. Chest. Med. 1 (2), 231–241.

Hammond, R. J., Baron, V. O., Oravcova, K., Lipworth, S., Gillespie, S. H., 2015. Phenotypic resistance in mycobacteria: is it because I am old or fat that I resist you? J. Antimicrob. Chemoth. 70 (10), 2823–2827.

Hendon-Dunn, C. L., Doris, K. S., Thomas, S. R., Allnutt, J. C., Marriott, A. A. N., Hatch, K. A., Watson, R. J., Bottley, G., Marsh, P. D., Taylor, S. C., et al., 2016. A flow cytometry method for rapidly assessing m. tuberculosis responses to antibiotics with different modes of action. Antimicrob. Agents Chemother., AAC–02712.

Hlatky, L., Alpen, E., 1985. Two-dimensional diffusion limited system for cell growth. Cell Prolif. 18 (6), 597–611.

Hu, Y., Mangan, J. A., Dhillon, J., Sole, K. M., Mitchison, D. A., Butcher, P. D., Coates, A. R., 2000. Detection of mrna transcripts and active transcription in persistent mycobacterium tuberculosisinduced by exposure to rifampin or pyrazinamide. Journal of bacteriology 182 (22), 6358–6365.

Keren, I., Kaldalu, N., Spoering, A., Wang, Y., Lewis, K., 2004. Persister cells and tolerance to antimicrobials. FEMS microbiology letters 230 (1), 13–18.

Keren, I., Minami, S., Rubin, E., Lewis, K., 2011. Characterization and transcriptome analysis of mycobacterium tuberculosis persisters. MBio 2 (3), e00100–11.

Kjellsson, M. C., Via, L. E., Goh, A., Weiner, D., Low, K. M., Kern, S., Pillai, G., Barry, C. E., Dartois, V., 2012. Pharmacokinetic evaluation of the penetration of antituberculosis agents in rabbit pulmonary lesions. Antimicrobial agents and chemotherapy 56 (1), 446–457.

Krombach, F., Mu¨nzing, S., Allmeling, A.-M., Gerlach, J. T., Behr, J., Do¨rger, M., 1997. Cell size of alveolar macrophages: an interspecies comparison. Environ. Health Perspect. 105 (Suppl 5), 1261.

Lipworth, S., Hammond, R., Baron, V., Hu, Y., Coates, A., Gillespie, S., 2016. Defining dormancy in mycobacterial disease. Tuberculosis 99, 131–142.

Macklin, P., Edgerton, M. E., Thompson, A. M., Cristini, V., 2012. Patient-calibrated agent-based modelling of ductal carcinoma in situ (dcis): from microscopic measurements to macroscopic predictions of clinical progression. J. Theor. Biol. 301, 122–140.

Manina, G., Dhar, N., McKinney, J. D., 2015. Stress and host immunity amplify mycobacterium tuberculosis phenotypic heterogeneity and induce nongrowing metabolically active forms. Cell host & microbe 17 (1), 32–46.

Marino, S., El-Kebir, M., Kirschner, D., 2011. A hybrid multi-compartment model of granuloma formation and T cell priming in tuberculosis. J. Theor. Biol. 280 (1), 50–62.

Matzavinos, A., Kao, C.-Y., Green, J. E. F., Sutradhar, A., Miller, M., Friedman, A., 2009. Modeling oxygen transport in surgical tissue transfer. Proc. Natl. Acad. Sci 106 (29), 12091–12096.

Osei, E., Akweongo, P., Binka, F., 2015. Factors associated with delay in diagnosis among tuberculosis patients in hohoe municipality, ghana. BMC Public Health 15 (1), 721.

Owen, M. R., Byrne, H. M., Lewis, C. E., 2004. Mathematical modelling of the use of macrophages as vehicles for drug delivery to hypoxic tumour sites. J. Theor. Biol. 226 (4), 377–391.

Patel, A. A., Gawlinski, E. T., Lemieux, S. K., Gatenby, R. A., 2001. A cellular automaton model of early tumor growth and invasion: the effects of native tissue vascularity and increased anaerobic tumor metabolism. J. Theor. Biol. 213 (3), 315–331.

Phillips, P. P., Mendel, C. M., Burger, D. A., Crook, A. M., Nunn, A. J., Dawson, R., Diacon, A. H., Gillespie, S. H., 2016. Limited role of culture conversion for decision-making in individual patient care and for advancing novel regimens to confirmatory clinical trials. BMC Med. 14 (1), 1.

Pienaar, E., Cilfone, N. A., Lin, P. L., Dartois, V., Mattila, J. T., Butler, J. R., Flynn, J. L., Kirschner, D. E., Linderman, J. J., 2015. A computational tool integrating host immunity with antibiotic dynamics to study tuberculosis treatment. J. Theor. Biol. 367, 166–179.

Pienaar, E., Matern, W. M., Linderman, J. J., Bader, J. S., Kirschner, D. E., 2016. Multiscale model of mycobacterium tuberculosis infection maps metabolite and gene perturbations to granuloma sterilization predictions. Infect. Immun. 84 (5), 1650–1669.

Powathil, G. G., Gordon, K. E., Hill, L. A., Chaplain, M. A., 2012. Modelling the effects of cell-cycle heterogeneity on the response of a solid tumour to chemotherapy: biological insights from a hybrid multiscale cellular automaton model. J. Theor. Biol. 308, 1–19.

Prideaux, B., Via, L. E., Zimmerman, M. D., Eum, S., Sarathy, J., O’Brien, P., Chen, C., Kaya, F., Weiner, D. M., Chen, P.-Y., et al., 2015. The association between sterilizing activity and drug distribution into tuberculosis lesions. Nat. Med. 21 (10), 1223–1227.

Sarathy, J. P., Via, L. E., Weiner, D., Blanc, L., Boshoff, H., Eugenin, E. A., Barry, C. E., Dartois, V. A., 2018. Extreme drug tolerance of mycobacterium tuberculosis in caseum. Antimicrobial agents and chemotherapy 62 (2), e02266–17.

Segovia-Juarez, J. L., Ganguli, S., Kirschner, D., Dec 2004. Identifying control mechanisms of granuloma formation during M. tuberculosis infection using an agent-based model. J. Theor. Biol. 231 (3), 357–376.

Sershen, C. L., Plimpton, S. J., May, E. E., 2016. Oxygen modulates the effectiveness of granuloma mediated host response to mycobacterium tuberculosis: a multiscale computational biology approach. Frontiers in cellular and infection microbiology 6.

Shorten, R., McGregor, A., Platt, S., Jenkins, C., Lipman, M., Gillespie, S., Charalambous, B., McHugh, T., 2013. When is an outbreak not an outbreak? fit, divergent strains of mycobacterium tuberculosis display independent evolution of drug resistance in a large london outbreak. J. Antimicrob. Chemoth. 68 (3), 543–549.

Singapore, T. S. M. R. C., 1981. Clinical trial of six-month and four-month regimens of chemotherapy in the treatment of pulmonary tuberculosis: The results up to 30 months. Tubercle 62 (2), 95–102. URL http://www.sciencedirect.com/science/article/pii/0041387981900167

Sprent, J., 1993. Lifespans of naive, memory and effector lymphocytes. Current opinion in immunology 5 (3), 433–438.

Study, E. A. M. R. C., 1981. Controlled clinical trial of five short-course (4-month) chemotherapy regimens in pulmonary tuberculosis: Second report of the 4th study. American Review of Respiratory Disease 123 (2), 165–170.

Swat, M. H., Thomas, G. L., Belmonte, J. M., Shirinifard, A., Hmeljak, D., Glazier, J. A., 2012. Multi-scale modeling of tissues using compucell3d. Methods Cell Biol. 110, 325.

Van Furth, R., Diesselhoff-den Dulk, M. M., Mattie, H., 1973. Quantitative study on the production and kinetics of mononuclear phagocytes during an acute inflammatory reaction. J. Exp. Med. 138 (6), 1314–1330.

Via, L. E., Savic, R., Weiner, D. M., Zimmerman, M. D., Prideaux, B., Irwin, S. M., Lyon, E., O’Brien, P., Gopal, P., Eum, S., et al., 2015. Host-mediated bioactivation of pyrazinamide: Implications for efficacy, resistance, and therapeutic alternatives. ACS Infect. Dis. 1 (5), 203–214.

Walz, A., Kunkel, S. L., Strieter, R. M., 1996. Cxc chemokines–an overview. Chemokines in Disease. RG Landes, Austin 1.

Wayne, L. G., Hayes, L. G., 1996. An in vitro model for sequential study of shiftdown of mycobacterium tuberculosis through two stages of nonreplicating persistence. Infection and immunity 64 (6), 2062–2069.

Wayne, L. G., Sramek, H. A., 1994. Metronidazole is bactericidal to dormant cells of mycobacterium tuberculosis. Antimicrobial agents and chemotherapy 38 (9), 2054–2058.

Zhang, L., Wang, Z., Sagotsky, J. A., Deisboeck, T. S., 2009. Multiscale agent-based cancer modeling. J. Math. Biol. 58 (4-5), 545–559.

